# Reproductive isolation of hybrid populations driven by genetic incompatibilities

**DOI:** 10.1101/007518

**Authors:** Molly Schumer, Rongfeng Cui, Gil G. Rosenthal, Peter Andolfatto

## Abstract

Despite its role in homogenizing populations, hybridization has also been proposed as a means to generate new species. The conceptual basis for this idea is that hybridization can result in novel phenotypes through recombination between the parental genomes, allowing a hybrid population to occupy ecological niches unavailable to parental species. A key feature of these models is that these novel phenotypes ecologically isolate hybrid populations from parental populations, precipitating speciation. Here we present an alternative model of the evolution of reproductive isolation in hybrid populations that occurs as a simple consequence of selection against incompatibilities. Unlike previous models, our model does not require small population sizes, the availability of new niches for hybrids or ecological or sexual selection on hybrid traits. We show that reproductive isolation between hybrids and parents evolves frequently and rapidly under this model, even in the presence of substantial ongoing migration with parental species and strong selection against hybrids. Our model predicts that multiple distinct hybrid species can emerge from replicate hybrid populations formed from the same parental species, potentially generating patterns of species diversity and relatedness that resemble an adaptive radiation.

## Introduction

The evolutionary significance of hybridization has been a hotly debated topic for decades (1). Homoploid hybrid speciation, speciation that occurs as a result of hybridization without a ploidy change (2), is generally thought to be an exceptionally rare outcome of hybridization and there are only a handful of well-supported cases of this phenomenon (3). Though it is not uncommon for species’ genomes to exhibit evidence of past hybridization, hybrids are often thought to be weakly isolated from parental species, though few studies have explicitly investigated this. The most convincing cases of homoploid hybrid speciation to date have identified hybrid populations that have transgressive traits that differentiate their ecologically or sexually selected phenotypes from parental species, establishing reproductive isolation between hybrids and parents (4–7).

Partly as an outgrowth of this empirical work, recent theoretical work has focused on transgressive traits generated via hybridization and their role in facilitating reproductive isolation between parental and hybrid populations. These models have shown that hybrid speciation is a likely outcome if a new niche is available and transgressive traits allow hybrids to colonize that niche (8, 9; discussed in 10, 11). However, it is generally difficult to evaluate the likelihood of generating novel traits via hybridization (but see 12) or the likelihood that the trait will allow hybrids to colonize a new environment, making it challenging to predict the importance of hybridization in generating new species. Other models of homoploid hybrid speciation have considered the potential for combined effects of inbreeding and chromosomal rearrangements to generate reproductively isolated hybrid species (13, 14), and empirical results from sunflowers provide support for this mechanism (15).

We present a different but simple model in which reproductive isolation between hybrid and parental populations emerges as a consequence of selection on genetic incompatibilities in a hybrid swarm. The first genetic model of speciation, described by Bateson, Dobzhansky and Muller (the BDM model, Fig. S1, 16–18), predicts that mutations at two genetic loci differentially accumulating along two lineages can negatively interact in their hybrids. The genetic incompatibility of hybrids constitutes a key component of reproductive isolation between many species, and is the basis for the biological species concept (18). Empirical research has suggested that these types of interactions are remarkably common (19–23; reviewed in 18, 24, 25). Though the theory of BDM incompatibilities was originally formulated in the context of allopatrically diverging species, our model investigates the fate of these incompatibilities in hybrid populations.

Under the simplest BDM scenario, derived genotypes are neutral, meaning that they have the same fitness as ancestral genotypes. Following hybridization, selection on these incompatibilities in a hybrid swarm is expected to result in the fixation of genotype combinations that are not incompatible with either parental species. However, incompatibilities may also frequently arise due to adaptive evolution within lineages or as coevolving pairs of loci (Fig. S1, S2, 18, 26–28). Here we show that selection on these types of incompatibility pairs can result in the fixation of genotype combinations that contribute to isolation between the hybrid population and one or the other parental species. In the presence of multiple pairs of such incompatibilities, this process can result in the rapid evolution of reproductive isolation of hybrid populations from both parental species.

Two features of this model make it particularly plausible biologically. First, empirical studies have revealed that hybrid incompatibilities rapidly accumulate between species (24), providing ample raw material for our model to work. Second, hybrid populations in which hybrids outnumber parental species occur frequently in nature, either in stable hybrid zones (29, 30) or as a result of environmental disturbance (31, 32). These characteristics suggest that the evolution of reproductive isolation of hybrid populations by this mechanism, a first step towards homoploid hybrid speciation, could be a common evolutionary outcome of hybridization.

## Results

### Modeling selection against hybrid incompatibilities

In the simplest model of a hybrid population, a mixture of individuals from both parental species form a new isolated population and mate randomly with respect to genotype (Fig. 1A). Using theory developed by Karlin (33), one can model the effect of selection at two polymorphic loci on gamete frequencies of a diploid sexual population (see Methods and Supplement 1). Using this two-locus selection model, one can see that the direction of fixation depends on the initial frequency of the parental alleles (*f*, see Fig. 1A and S3) and dominance at each locus (*h*, see Fig. S3).

**Fig. 1.**
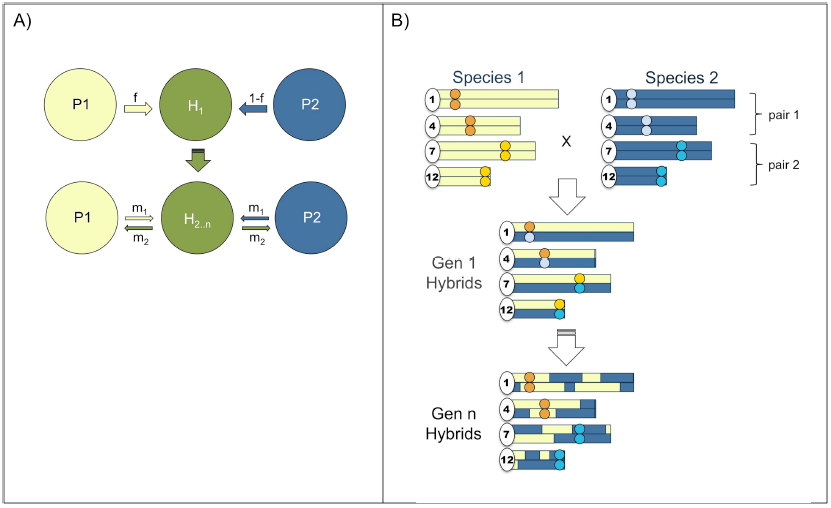
Schematic of the simplest hybrid speciation scenario. The simplest model of hybrid speciation evolves in a hybrid swarm (A) via fixation of parental genetic incompatibility pairs in opposite directions (B). *f* is the proportion of the hybrid (H) population colonized by parent 1 (P1), m_1,2_ denotes migration rates between the parental and hybrid populations over n generations.

This purely deterministic model of selection on hybrid incompatibilities is unrealistic because even large populations experience some degree of genetic drift. We thus extended the model to include genetic drift, which can affect the speed and direction of fixation of incompatibility pairs (Fig. S4). For coevolving or adaptive BDM incompatibilities (Supporting Information 1, Fig. S1, S2), this model predicts that at equal admixture proportions (*f* = 0.5), a single codominant incompatibility pair has a 50% chance of fixing for either parental allele combination (Fig. 2, Supporting Information 2). Interestingly, while genetic drift in very small populations could accomplish the same thing (8), the process described here occurs rapidly in large populations and is driven by deterministic selection (Fig. 2). Given this result, it is clear that large hybrid populations with two or more hybrid incompatibilities could, in principal, fix for one parental genotype at one incompatibility pair and the other parental genotype at the other incompatibility pair (Fig. 1B). This outcome would result in reproductive isolation of the hybrid population from both parental species.

**Fig. 2.**
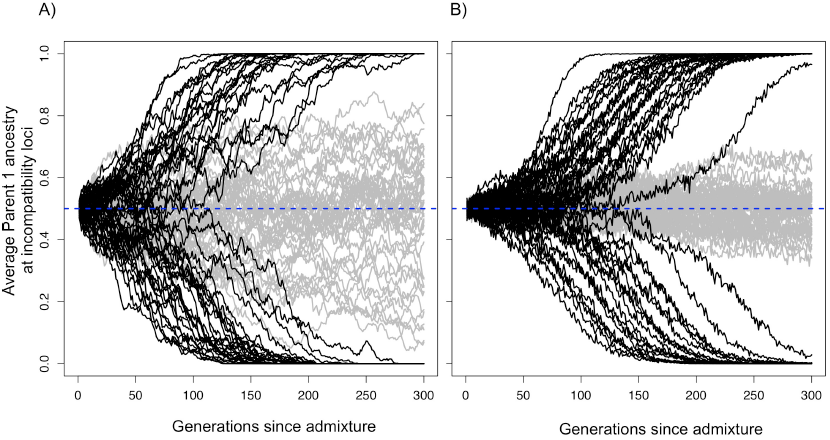
Hybrid populations rapidly fix for hybrid incompatibility locus pairs. Selection drives hybrid incompatibility loci to fixation, even when a hybrid population forms at equal admixture proportions (*f=*0.5). Black lines show average parent 1 ancestry at a hybrid incompatibility pair (*h*=0.5, s=0.1) in a population of (A) N=1,000 or (B) N=10,000 diploid individuals. Gray lines show results for this same population size with no selection. Because of this behavior, when two incompatibility pairs exist they may fix for opposite parents, resulting in reproductive isolation of hybrids from both parents (see Fig. 1B).

### Simulations of an isolated hybrid population

To investigate the dynamics of multiple incompatibility pairs, we simulated a large, randomly-mating and spatially isolated hybrid population with two pairs of unlinked hybrid incompatibility loci (see Methods; Fig. 1A, setting *m_1_*=*m_2_*=0). The fitness scheme used is that of a coevolutionary incompatibility model (Fig. S2), assuming that incompatibilities are codominant (i.e. *h*=0.5), that fitness is symmetric with respect to the parental source of alleles (i.e. *w_ij_* = *w_ji_*) and that the cumulative fitness effects of multiple incompatibility pairs is multiplicative. If hybrid populations fixed for the parent 1 genotypes at one incompatibility pair and the parent 2 genotypes at the other, we considered the hybrid population as having evolved reproductive isolation from both parents.

In simulations of this simple scenario, reproductive isolation between hybrid and parental populations evolved frequently and rapidly. For two incompatibility pairs with selection coefficients (*s*) of 0.1, 47±2% of simulated hybrid populations became isolated from both parental species within an average of ∼150 generations. Exploring a range of *s* (0.1–0.5, Fig. S5), initial admixture proportions (*f*=0.3–0.7, Fig. S6), and population sizes (100-10,000 diploids), we conclude that, unless fitness of hybrids is close to 0 or ancestry is highly skewed, reproductive isolation evolves rapidly and with surprisingly high probability (27±2% to 44±2% of the time; on average within 55 ± 14 to 190 ± 43 generations, see Supporting Information 3).

### The effect of dominance and asymmetry in selection intensity

In the above simulations, we assume that selection on different hybrid incompatibility interactions is symmetrical (*s*_1_=*s*_2_, Fig. S2), but it is unlikely that selection is truly equal on different hybrid genotype combinations. When fitness is completely asymmetrical (i.e. *s*_1_ or *s*_2_ =0, as for neutral BDM incompatibilities), only strong genetic drift can cause the fixation of genotype pairs that are incompatible with either parent (Fig. S8, Supporting Information 4) as selection cannot do so (see Fig. S1, S3, S7, Supporting Information 4). This reliance on genetic drift implies that this process will be slow unless an extreme bottleneck is invoked.

In contrast, the dynamics of BDM incompatibilities resulting from adaptation within parental lineages can be quite different (Fig. S1). Notably, while selection may also be highly asymmetric in such cases (34), derived alleles have higher fitness than ancestral alleles resulting in the fixation of genotype combinations that are incompatible with both parents. We find that isolation evolves with similar frequency under asymmetric selection as long as selection is strong relative to drift (Supporting Information 3D), because even weak selection will prevent the fixation of the ancestral genotype.

We arbitrarily simulate codominant hybrid incompatibilities, but Karlin’s model (Fig. S3) shows that patterns of fixation are different depending on the value of *h*. In particular, when *h* is zero or unity, fixation is not strongly dependent on admixture proportions (Fig. S3). When we simulate variation in dominance among incompatibility interactions (see Supporting Information 3E), we find that reproductive isolation between hybrid populations and parental species evolves with comparable frequency (44–48±2% vs 47±2% under the codominant scenario).

### Increasing the number of hybrid incompatibilities

Recent empirical studies have suggested that most species are distinguished by multiple hybrid incompatibilities (24, 35–40). Theoretically, barring extinction of the hybrid population, increasing the number of incompatibilities should increase the probability that a hybrid population will evolve isolation from both parental species. In order to illustrate this, we simulated 3–6 unlinked hybrid incompatibility pairs (Supporting Information 5). As expected, increasing the number of hybrid incompatibilities increases the probability that the hybrid population will be isolated from each parent by at least one incompatibility (>96% with 6 incompatibility pairs, Fig. 4, Fig. S5, Supporting Information 5B).

**Fig. 3.**
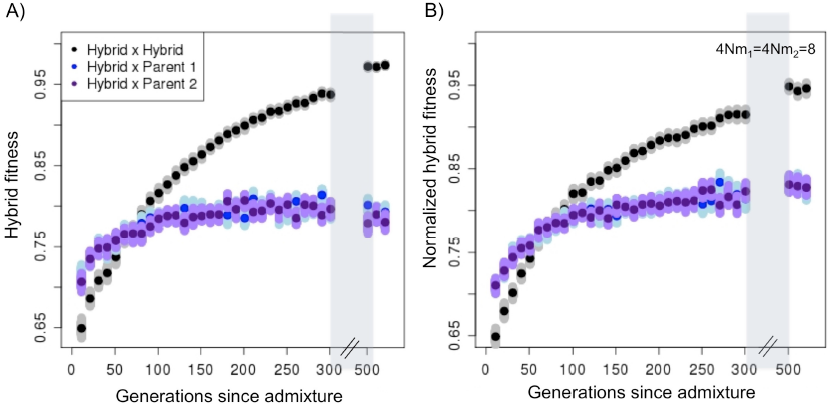
Hybrid populations rapidly develop reproductive isolation from both parental species, even in the presence of migration. (A) Once hybrid populations diverge in ancestry at hybrid incompatibility loci from parental populations, individual hybrids have higher fitness on average when they mate with other hybrids in their population compared to either parent (B) even with ongoing migration (4*Nm*_1_=4*Nm*_2_=8) from parental populations. Dark point indicates mean fitness, smears indicate 1,000 bootstrap resamplings of the mean. Simulation parameters: 100 replicates per time point, N=1000, 20 hybrid incompatibility pairs, *s_1_, s_2_* and *h* drawn from a distribution (see details in Supporting Information 5D).

**Fig. 4.**
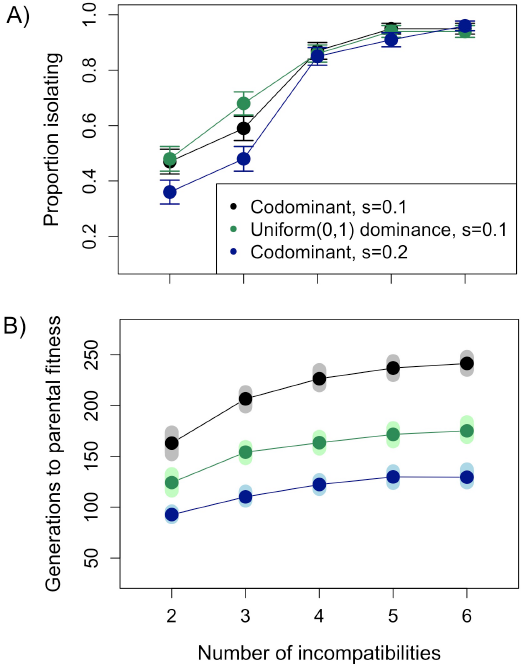
Relationship between the number of hybrid incompatibility pairs and probability of evolving isolation from both parents. With an increasing number of hybrid incompatibility pairs, reproductive isolation from both parents increases in likelihood (A) but populations require longer periods of time to reach parental fitness levels (B). In these simulations two to six hybrid incompatibility pairs distinguish the hybridizing species (*s*=0.1) and hybrid populations formed at equal admixture proportions (*f*=0.5, 1000 diploid individuals). In (A) error bars show two standard deviations of the mean; in (B) smears show 1000 bootstrap resamplings of the mean. Results based on 500 simulations.

We assume in most of our simulations that loci involved in hybrid incompatibilities are completely unlinked. As the number of incompatibilities increases, this becomes increasingly less likely. Physical linkage between loci involved in different epistatic interactions can reduce the frequency at which hybrid populations evolve isolation because sites are more likely to fix for the same parent (Fig. S9, Supporting Information 5C). Interestingly, when hybrid incompatibility loci are linked to a neutral BDM incompatibility, this does not significantly lower the frequency at which hybrid populations evolve reproductive isolation (Supporting Information 5B). Furthermore, linkage between coevolving incompatibilities and neutral BDM incompatibilities can more frequently result in fixation of neutral BDM incompatibilities for a parental genotype (21±2%), resulting in stronger isolation between hybrid and parental populations (Supporting Information 5B).

The above simulations focus on simple models of hybrid incompatibility that show this process can occur in principle. To capture more biological realism in the number and types of incompatibilities, we simulated 20 incompatibility pairs with randomly determined genomic position and dominance, drawing selection coefficients from an exponential distribution with a mean *s*=0.05 and introducing asymmetry in selection (see Supporting Information 5D). In these simulations 95% of populations developed isolation from both parents within 500 generations. On average, hybrid population first developed isolation from both parents after 180 generations and were isolated by 7 incompatibility pairs by 500 generations. Interestingly, because incompatibilities with the largest fitness effects tend to fix first, hybrid populations developed considerable reproductive isolation from parental species even before all incompatibilities were fixed in the population (Fig. 3, Fig. S10). Overall, our simulations suggest that rapid evolution of reproductive isolation of hybrid populations is likely when parental species are separated by several hybrid incompatibilities with varying levels of asymmetry, linkage and dominance.

### Simulations of hybrid populations with migration

We model hybrid population formation as a discrete event, but many hybrid populations experience continued gene flow with parental species. This process may impede the evolution of reproductive isolation by preventing the fixation of genetic incompatibilities. To evaluate this, we simulated hybridization scenarios with ongoing migration (Fig. 1A, 4*Nm*=87–40). Even with substantial continuing gene flow from parental populations, hybrid populations evolved reproductive isolation from them at high frequencies (i.e. 37±2% of simulations with two incompatibilities, *s*=0.1 and 4*Nm*=8; Supporting Information 6).

Intuitively, as chance plays an important role in which incompatibility pairs fix, independently formed hybrid populations from the same parental species can evolve isolation from each other. To demonstrate this effect, we simulated two hybrid populations formed from the same parental species (Fig. S12). In the absence of migration, the two hybrid populations evolved isolation from each other frequently (52±5%, as expected given two hybrid incompatibilities, see Supporting Information 6C). Remarkably, this outcome was still observed frequently with relatively high gene flow between the two hybrid populations (18±4% with 4*Nm*=8 and two hybrid incompatibilities, Supporting Information 6C).

## Discussion

We describe a new model of the evolution of reproductive isolation of hybrid populations, a first step towards hybrid speciation. Unlike previous models of hybrid speciation, our model does not rely on transgressive hybrid traits, the availability of new ecological niches, or small population sizes, but rather deterministic selection in large populations against hybrid incompatibilities. Reproductive isolation can evolve rapidly under our proposed model; with moderate selection (i.e. *s*=0.1) on a small number incompatibility pairs in an allopatric hybrid population, reproductive barriers from both parents emerge with close to 50% probability and on average within ∼150 generations. Hybrid reproductive isolation also evolves frequently with substantial levels of ongoing migration between hybrids and parental species. The widespread existence of hybrid incompatibilities and the rapid emergence of reproductive isolation predicted by this model imply that hybrid speciation by this mechanism could be a common natural consequence of hybridization.

It is important to note several factors that may influence how common this mode of hybrid reproductive isolation is in natural populations. In our model, isolation between hybrids and parents is inherently weaker than between the two parents. Nonetheless, we propose that this mechanism may be a crucial first step in limiting gene flow between hybrids and parents, allowing for the development of other isolating mechanisms. For example, theoretical work predicts that reinforcement can develop even when selection is moderate on hybrids (e.g. s∼0.3, 41–44). We also note that our model only represents fitness in terms of genetic incompatibilities and that hybrid populations can have lower fitness as a result of ecological or sexual selection. For example, when parental species exert negative sexual selection against hybrids, hybrid populations are significantly more likely to be outcompeted by parents (Supporting Information 6C). This highlights the fact that the plausibility of this model depends on the biology of the species in question; parental species of swordtail fish and pupfish do not discriminate strongly against hybrids (45–47), while this is not the case in mice (48).

An additional consideration is that this model will only be relevant during a particular window of divergence between species. When the fitness of hybrid populations is low, they are more prone to extinction or displacement by parental populations (Supporting Information 3). This suggests that hybrid speciation is most likely to occur in a period of evolutionary divergence during which species have accumulated some hybrid incompatibilities but have not diverged to the point to which hybrids are largely inviable. The most detailed work characterizing genetic incompatibilities has been in *Drosophila* species, where hybrids generally have substantially reduced fitness compared to parents (36, 37, 49). Hybrids between several other species studied to date, however, are affected by fewer incompatibilities or incompatibilities of weaker effects (20, 35, 40, 50–53). Such species could be more likely to form hybrid populations, and should be the focus of future empirical research.

It is also important to note that reduced frequency of reproductive isolation with increasing selection on hybrids can be mediated by an increase in the total number of hybrid incompatibilities. In our simulations, we see a positive relationship between the number of interactions and the probability of developing reproductive isolation (Fig. 4) and a negative relationship between the total strength of selection on hybrids and the probability of developing reproductive isolation (Fig. S5). This tradeoff suggests that reproductive isolation can evolve between hybrid and parental populations even when the fitness of hybrids is low (as in Fig. 3, Fig. S5).

Our model requires that coevolving incompatibilities or BDM incompatibilities arising from adaptive evolution frequently occur between species. Accumulating evidence suggests that incompatibilities arising from coevolution may be common (24, 27, 54–57). For example, in marine copepods, coevolution between cytochrome *c* and cytochrome *c* oxidase results in a reciprocal breakdown of protein function in hybrids (57). In addition, the fact that many known incompatibility genes involve sexual conflict, selfish genetic elements, or pathogen defense suggests an important role for coevolution in the origin of incompatibilities (27, 54, 58, 59). Our model also applies to BDM incompatibilities that arise due to within-lineage adaptation. It is currently unknown whether effectively neutral BDMIs are more common than adaptive BDMIs. Though many incompatibilities have asymmetric selection on different hybrid genotypes (e.g. 22, 23, 60), neutrality has not been established in these cases. Anecdotal evidence supports the idea that adaptive incompatibilities are common since many hybrid incompatibilities show evidence of positive selection within lineages (61), but the relative frequency of adaptive and neutral BDMIs awaits answers from further empirical research.

The patterns predicted by this model are testable with empirical approaches. A large number of studies have successfully mapped genetic incompatibilities distinguishing species (19, 20, 36, 37, 39, 53, 62). Ancestry at these sites can be determined in putative hybrid species, and the relative contribution of parental-derived incompatibilities to reproductive isolation can be determined experimentally. For some species, it may be possible to evaluate the dynamics of incompatibilities relative to the genetic background in hybrid swarms (63). We predict that many hybrid populations exhibiting postzygotic isolation from parental species will have fixed incompatibility pairs for each parental species. Some proposed cases of hybrid speciation report reduced fitness of offspring between parental and hybrid species consistent with the mechanism described here (5, 15, 64, 65) and are promising cases for further empirical research.

An intriguing implication of our model is that independently formed hybrid populations between the same parental species can develop reproductive isolation from each other. The likelihood of this outcome increases with the number of incompatibilities. In sunflowers, empirical studies of ecologically-mediated hybrid speciation have identified multiple hybrid species derived from the same parental species (66). It is interesting to note that selection against hybrid incompatibilities could generate the same pattern in replicate hybrid populations. In fact, this mechanism could generate a species diversity pattern similar to that expected from an adaptive radiation, with multiple closely related species arising in a short evolutionary window. This finding is striking because our model does not invoke adaptation and suggests that non-adaptive processes (i.e. selection against incompatibilities) could also explain clusters of rapidly arising, closely related species.

## Methods

### Mathematical model of selection on hybrid incompatibilities

To characterize evolution at hybrid incompatibility loci in hybrid populations without drift, we used the equations described by Karlin (33) to calculate changes in allele frequency as a result of two-locus selection. The frequency of gamete *i* at generation *t* is given by

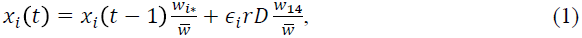

where ϵ_1_= ϵ_4_= -ϵ_2_-ϵ_3_= -1 and the marginal fitness of allele *i*,

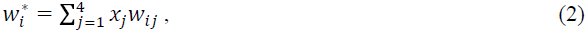

the mean fitness of the population,

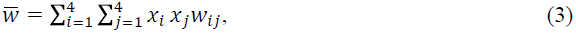

*W*_14_ is the fitness of a double heterozygote, *r* is the recombination rate and *D* is linkage disequilibrium between the two loci. These equations assume random mating, non-overlapping generations and that fitness depends only on two-locus genotype and not on whether the chromosome was maternally or paternally inherited (*w_ij_*=*w_ji_*). To model changes in allele frequencies over time we developed a custom R script (available from Dryad, doi:XXX). Iterating through the change in allele frequencies each generation as a result of selection gives the expected patterns of fixation at incompatibility loci without genetic drift (Fig. S3).

Karlin’s model of fixation of hybrid incompatibilities does not realistically predict expected patterns in natural populations because even large populations will have some level of genetic drift. To model this we added multinomial sampling of N diploid individuals and recalculated allele frequencies each generation (available from Dryad, doi:XXX). Patterns of fixation incorporating genetic drift through multinomial sampling show similar dynamics to the model lacking genetic drift, with the exception of several equilibrium states specific to the latter (see Fig. S4, Supporting Information 2).

### Description of simulation program

In order to simulate multiple pairs of hybrid incompatibilities in a large number of individuals and demographic scenarios, we developed a custom c++ program, called `admix’em` (github: https://github.com/melop/admixem). The code allows one to specify the number and length of chromosomes and the genomic locations of hybrid incompatibilities and neutral markers. The current implementation assumes non-overlapping generations and diploid sexual individuals. When modeling linkage, we assume a uniform recombination rate and one recombination event per chromosome per meiosis. Unless otherwise specified, we model all pairs of hybrid incompatibility loci as unlinked and on separate chromosomes. As we are interested in short-term dynamics, we do not implement mutation.

Selection coefficients are assigned to particular allelic combinations according to a hybrid fitness matrix (see Fig. S1,S2). Based on each individual’s genotype at hybrid incompatibility loci, we calculate total individual fitness *w*, defined as the probability of survival of that individual. Total fitness across multiple incompatibility pairs is assumed to be multiplicative. Each female mates with one randomly selected male (but incorporate assortative mating in Supporting Information 6C), and produces a Poisson distributed number of offspring with a mean of 2. After selection, if the carrying capacity (N) is not reached, additional offspring from the same mating events will be drawn from a Poisson distribution with a new mean of (carrying capacity – current population size)/number of females. This process is repeated until carrying capacity is reached or females have no available gametes (set to a maximum of 10).

All reported results are based on 500 replicate simulations, which were conducted for 2000 generations. In the majority of simulations (except Supporting Information 5B,C) the hybrid population is initially colonized by 500 individuals of each parental species. Hybrid and parental populations were modeled as spatially distinct with migration parameters between them; most simulations specified one hybrid population formed between two parental populations (Fig. 1A) but we also simulated a stepping-stone model and a model with multiple independently formed hybrid populations (Supporting Information 6, Fig. S11, Fig. S12). Details on individual simulations and results can be found in the supporting text. Hybrid populations are considered to have evolved reproductive barriers from both parents if they fix at least one incompatibility for each parent; the actual strength of selection against offspring between hybrids and parents will depend on the selection coefficient and number of incompatibilities.

## Acknowledgements

We would like to thank members of the Rosenthal and Andolfatto labs for helpful discussion and Clair Han, Ying Zhen, and Yaniv Brandvain for comments on earlier versions of this manuscript. This work was supported by an NSF GRFP (DGE0646086) and NSF DDIG (DEB-1405232) to M.S and an NSF IOS-0923825 to GGR.

## Author contributions

M.S. conceived of the project. M.S., R.C., G.G.R, and P.A. designed the project. M.S., R.C and P.A. wrote scripts and ran simulations. R.C. wrote admix’em program. M.S., R.C., G.G.R and P.A. analyzed results and wrote the manuscript.

